# Increased force and elastic energy storage are not the mechanisms that improve jump performance with accentuated eccentric loading during a constrained vertical jump

**DOI:** 10.1101/2023.10.30.564851

**Authors:** Eric Yung-Sheng Su, Timothy J. Carroll, Dominic J. Farris, Glen Lichtwark

## Abstract

**Objective:** Accentuated eccentric loading (AEL) involves higher load applied during the eccentric phase of a stretch-shortening cycle movement, followed by a sudden removal of load before the concentric phase. Previous studies suggest that AEL enhances human countermovement jump performance, however the mechanism is not fully understood. Here we explore whether isolating additional load during the countermovement is sufficient to increase ground reaction force, and hence elastic energy stored, at the start of the upward movement and whether this leads to increased jump height or power generation.

**Methods:** We conducted a trunk-constrained vertical jump test on a custom-built device to isolate the effect of additional load while controlling for effects of squat depth, arm swing, and coordination. Twelve healthy, recreationally active adults (7 males, 5 females) performed maximal jumps without AEL, followed by randomised AEL conditions prescribed as a percentage of body mass (10%, 20%, and 30%), before repeating jumps without AEL. **Results.** No significant changes in vertical ground reaction force at the turning point were observed. High load AEL conditions (20% and 30% body weight) led to slight reductions in jump height, primarily due to decreased hip joint and centre of mass work. AEL conditions did not alter peak or integrated activation levels of the knee extensor muscles.

**Conclusion:** These findings suggest that increased elastic energy return may not be the primary mechanism behind the potentiating effects of AEL on jump performance, and other factors such as rate of descent, squat depth, or body configuration may contribute to effective AEL.

## Introduction

Accentuated eccentric loading (AEL) is a form of movement manipulation that has been suggested to enhance power output during stretch-shortening cycle exercises. For example, during a human countermovement jump (CMJ), AEL requires an external load to be added to the body during the downward (eccentric) movement and then released at the transition from the downward to the upward (concentric) movement. Studies have indicated that the immediate response to AEL during a CMJ could increase jump height by ∼4.3-9.5%, increase peak power output by ∼9.4-23.2%, and increase maximal concentric vertical ground reaction force by ∼3.9-6.3%. ^1,2^ By contrast, Aboodarda et al. ^3^ reported that AEL applied through elastic resistance in human drop jumps did not alter jump height, muscle activation level, or other kinetic profiles during push-off. There is conflicting evidence on the performance-enhancing effect of AEL CMJ, with studies showing both positive ^1,2,4^ and no effect. ^3,5–7^ A review study concluded that current evidence for acute responses to AEL is inconsistent, possibly due to different exercises selected, equipment used, range of loads, or participants’ characteristics across different experiments. ^8^

There are a number of potential mechanisms that might drive enhanced power output during AEL movements. The most common explanation for why AEL should enhance power is that increased load in the eccentric phase amplifies elastic energy storage in the tendon and aponeurosis, which can be released in the concentric phase. ^9^ For instance, AEL CMJ may result in greater force generation in the descent to decelerate the added mass or resist the added force, potentially resulting in greater tendon loading and strain energy storage prior to the upward motion. Alternatively, AEL might change the coordination strategy, allowing muscles to generate greater force or power during the subsequent concentric portion of the jump. This might be achieved through increased squat depth ^10,11^ or changes in the position of the centre of mass (COM) relative to the joints. ^12^ Whilst there is some evidence that humans can achieve greater jump heights through AEL without significant changes in squat depth, ^2^ a combination of the factors above might contribute to enhanced jump performance in AEL.

Here we sought to investigate whether isolating the effect of additional load (in the absense of other potential contributors) during an AEL CMJ can increase the force applied to the ground at the start of the concentric movement in a CMJ and enhance positive power generation during the jump. In order to isolate whether additional load leads to increased force and therefore increased elastic energy storage, we used a constrained AEL scenario to control multiple alternative factors that might influence force and power. Specifically, we controlled squat depth, fore-aft and lateral movement of the trunk, arm swing and the point of application of the added mass during the CMJ. As such, the added load should be the dominant remaining factor to influence the force applied to the ground and hence elastic energy storage and return from lower limb tendinous tissues. We hypothesised that under these controlled conditions, AEL would increase the vertical ground reaction force (VGRF) at the start of the concentric movement and therefore increase the power and subsequent work during the concentric phase of the CMJ. We also hypothesised that the increased force would require greater muscle activity of the lower limb muscles undergoing stretch-shortening cycles, particularly the knee extensor muscles that are largely responsible for power production during the jump.

## Materials and Methods

### Sample size calculation

We conducted a pilot study with the similar experimental setup as Sheppard et al. ^2^ From our pilot study, we found an improvement in jump height with AEL with an effect size of 0.83 (paired analysis). Our sample size calculation found that 11 participants were sufficient to achieve a statistical power of 80% (within-factor effect size = 0.83, alpha = 0.05, one-tailed). We recruited one additional participant to account for potential data loss or improve the statistical power.

### Participant characteristics

Twelve healthy and recreationally active adults (7 males, 5 females, height = 177 ± 8.1 cm, mass = 75.3 ± 10.7 kg, age = 28.1 ± 6.7 years) gave written informed consent to participate in this study. Ethical approval was granted from the institutional ethics review committee at The University of Queensland (approval number: 2021/HE001129). Our study required participants to achieve an effective jump height of at least 40 cm (male) or 30 cm (female) using a vertical jump-and-reach device (Swift Performance, Wacol, QLD, Australia). The effective jump height definition was consistent with the Exercise and Sports Science Australia testing protocol. ^13^ Participants were screened for their jump ability prior to participation.

### Testing protocol

Participants attended two laboratory sessions, one for familiarization (session one) and the other for data collection (session two). In session one, participants performed 20-30 practice jumps on a jumping sled (see below – Jumping Sled) with and without AEL from various squat depths, with at least 1 minute rest between jumps. The sled only allowed the trunk to translate in the vertical direction, with no trunk rotation or fore-aft or lateral translation.

Participants practiced non-AEL and AEL jumps with 10%, 20%, and 30% additional body mass attached to the sled. The non-AEL condition included the mass of the backrest, ∼7 kg, and the other AEL conditions added an additional mass on top of the backrest mass.

Participants attempted maximal effort jumps with both hands holding onto bars located on the backrest above the shoulders, thus preventing arm swing. Based on the jumping performance in the non-AEL condition, we selected an appropriate squat depth for each participant that maximised jump height, which was used as the controlled squat depth for the remainder of the study. Participants practiced CMJ from this same depth across different AEL conditions. Squat depth was monitored live and recorded based on a string potentiometer attached to the sled. Visual feedbacks of squat depth and jump height were given after each jump.

In session two, we collected force plate, motion capture, and electromyography (EMG) data to examine jump performance under different AEL conditions. Participants performed 3∼5 submaximal non-AEL jumps to warm-up. They then performed at least 3 valid maximal jumps at their controlled squat depth (recorded in session one) for each of the different conditions. Participants were required to achieve the specified squat depth to qualify for a valid trial, with a maximum of 5 cm error margin below this depth. This was monitored using the string potentiometer with visual feedback given after each trial. Participants performed maximal jumps with non-AEL condition (BW_pre_) first, followed by AEL conditions (10%, 20%, and 30% AEL) in a randomised order, and finished with another non-AEL condition (BW_post_). At least 1 minute rest was provided between each maximal trial.

### Jumping Sled

The jumping sled design is shown in Figure 1. An aluminium backrest (sled) was attached to a squat cage fitted with linear rails and bearings. The sled included an electromagnet, shoulder bars, and a waist belt for participants. A pulley system with an inextensible metal wire and weight plates was used to add or remove extra mass. One end of the wire was attached to a light iron disc that would hold to the electromagnet when charged. A linear string potentiometer (SP2-50, TE Connectivity, Berwyn, PA, USA) was fixed on the squat cage to measure the sled’s vertical position. An electromagnetic switch was used to release the added mass when the participant reached the lowest depth during the jump. The mass was dropped safely into a box behind the participant.

**Figure 1.**
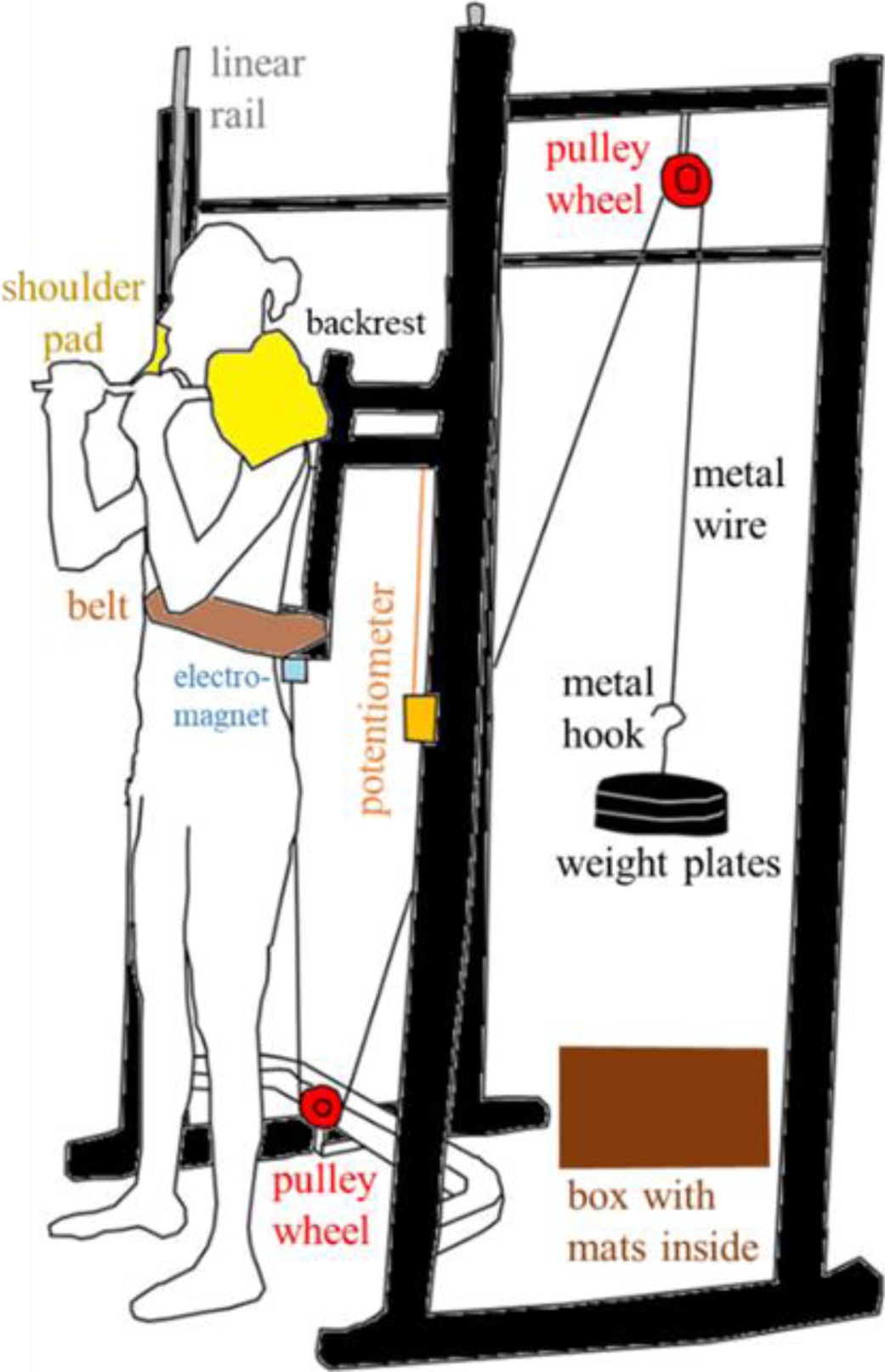
Jumping sled design. The squat cage (black) was fitted with linear rails (grey) and an aluminium backrest (black). An electromagnet (blue) was attached to the bottom of the backrest, and two shoulder bars with foam padding (yellow) were fixed at the top of the backrest. Participants were strapped firmly on the backrest with a waist belt (brown). Two pulley wheels (red) were fixed on the squat cage, and a metal wire was used to carry a metal hook and weight plates (black). A linear string potentiometer (orange) was attached between the fixed frame and the sliding backrest. The weight plates were released into a box (dark brown) behind the participant.

We recorded the calibrated potentiometer length in the A/D board (Micro 1401, Cambridge Electronic Device, Cambridge, United Kingdom) and ran a custom script to determine the release timing in the Spike 2 software (Cambridge Electronic Device, Cambridge, United Kingdom). We defined a threshold length equivalent to each participant’s controlled squat depth. The system monitored when the potentiometer velocity turned positive after crossing this threshold to ensure the additional mass was only released when the lowest depth was achieved. Once the participant reached the required depth and initiated the push-off phase, the Spike 2 software sent a TTL pulse to deactivate the electromagnet, causing the added mass to drop. On average, there was a delay of 0.084 ± 0.050 seconds from the turning point to TTL generation, during which the average torso COM displacement was 2.81 ± 1.84 cm (or 6.9 % of the average upwards jump displacement).

### Data collection

#### Motion Capture and Force Plates

Reflective markers (9mm) were placed on the participant’s skin and tracked with an 11- camera, 3D optoelectronic system (Oqus, Qualisys, AB, Sweden). These markers were placed on the following anatomical landmarks on the trunk and both legs: acromion process, jugular notch, xyphoid process, ASIS, PSIS, iliac crest directly superior to the greater trochanter, medial and lateral joint centre of the knee, medial and lateral malleoli, sustentaculum tali (medial calcaneus marker), fibular trochlea (lateral calcaneus marker), calcaneal tuberosity (posterior calcaneus marker), first and fifth metatarsal, interphalangeal joint of the big toe. A cluster of 4 markers were also attached to both the shanks and thighs to track motion. We also added one marker on each side of the deltoid muscle belly to allow tracking of the trunk segment motion. Marker positional data were sampled at 125 Hz and collected in Qualisys Track Manager (QTM) software (Qualisys, Gothenburg, Sweden). Two tri-axial AMTI in-ground force plates (AMTI, Watertown, MA, USA) synchronously collected ground reaction force (GRF) data at 1250 Hz in QTM. Prior to jumping trials, we collected a static trial for each participant, where they stood with legs evenly spaced on two force plates and their hands positioned similarly to the jumping trials.

#### Electromyography

Surface EMG data were collected from six lower limb muscles on the right leg with a wireless EMG system (MYON m320; MYON, Schwarzenberg, Switzerland). Muscles recorded were soleus (SOL), medial gastrocnemius (MG), vastus lateralis (VL), rectus femoris (RFEM), biceps femoris long head (BFEM), and gluteus maximus (GMAX). Bipolar EMG electrodes of 24 mm diameter were placed over the belly of each muscle according to the SENIAM guidelines. ^14^ EMG data were synchronously recorded with motion capture data in QTM at 1250 Hz and exported as MAT file for further analysis in MATLAB (Mathworks, Natick, MD, USA).

### Data Processing and Analysis

#### Kinematics and Kinetics

Motion capture markers were labelled, tracked, and digitally exported with EMG and GRF data. We used Opensim ^15,16^ and a publicly available model ^17^ that was first scaled to each participant and then performed inverse kinematics and inverse dynamics calculation in OpenSim. Scaling factors were calculated by dividing participant-specific marker distances by the corresponding distances on the generic model. Inverse kinematics results and experimental GRF data were filtered with a 15Hz low-pass second-order Butterworth filter prior to inverse dynamic calculation. Opensim outputs data were then imported into MATLAB for further analysis, however, we only analysed trials that achieved the highest effective jump height in each condition for each participant for kinematic and kinetic data.

In MATLAB, we calculated effective jump height as the difference between the highest vertical position of the manubrium marker (MAN) for each trial and the average MAN vertical position in the static trial for each participant. The start of the jump was defined as the instant when MAN vertical position fell below 1 cm from the average vertical position during the initial standing phase for each trial. The turning point of the jump was defined as the lowest vertical position of the MAN for each trial. Take-off was estimated when the right vertical GRF dropped to zero. Push-off phase was defined from the turning point to take-off. Squat depth was defined as the difference between the average MAN vertical position in the static trial for each participant and the lowest MAN vertical position for each trial. Average descent speed was calculated by dividing squat depth over the duration for descent. We took the dot product of the model’s joint moment and joint angular velocity to calculate joint power for each joint (right hip, knee, ankle) across all degrees of freedom. Positive values in joint moments represent hip extension, knee extension, and ankle plantarflexion internal moments. Vertical COM power during the concentric phase was calculated as the dot product of the model’s vertical COM velocity (from OpenSim Body Analysis) and the experimentally collected total vertical GRF data (sum of two vertical GRF from each plate). Joint moments, joint power, and vertical COM power data were filtered with a 5Hz low-pass second-order Butterworth filter. We then integrated joint power and vertical COM power during the push-off to calculate concentric joint work and concentric vertical COM work. Finally, we calculated the total lower limb joint work by summing the work across all degrees of freedom at the hip, knee, and ankle joints.

#### Electromyography

Raw EMG data were zero-phase band-pass filtered (30-350 Hz second-order Butterworth filter), rectified, and zero-phase low-pass filtered (5 Hz second-order Butterworth filter) to form a linear envelop as the processed EMG data. Due to an acquisition error, some trials in the AEL conditions had incomplete raw EMG data that only ranged from the start of the jump until just after the turning point. We excluded trials with incomplete EMG data, and reduced data to two key EMG outcome measures for each participant and condition: the peak EMG amplitude achieved during the entire jump and the integrated EMG during push-off phase.

These outcome measures (peak and integrated EMG) were calculated by averaging across multiple trials for each condition per participant. For each participant, the peak EMG amplitude obtained across all trials was used to normalise all EMG values, as per consensus recommendations. ^18^

### Statistical analysis

Linear mixed-effects models with Restricted Maximum Likelihood solution were used to compare AEL conditions (BW_pre_ [Non-AEL], 10%, 20%, 30% AEL). A mixed-effects model to perform repeated measures analysis was used to account for one missing data point in the 30% AEL condition. We reported the p-value and the partial eta squared (η^2^_p_) effect size for each main effect. Dunnett’s multiple comparison tests were used to detect the origin of significant main effects between experimental conditions (10%, 20%, 30% AEL) against one single control condition (BW_pre_). We reported the p-value and the Cohen’s d effect size for each multiple comparison test. Paired t-tests were used to compare non-AEL conditions before (BW_pre_) and after (BW_post_) AEL interventions, and the p-value and the Cohen’s d effect size were reported. Alpha was set at 0.05. Statistical assumptions for each dependent variable were performed using the D’Agostino-Pearson normality test and all data were normally distributed. Statistical analysis was performed using GraphPad Prism 8.

## Results

### Jump Height and Rate of Descent

There was a significant main effect of AEL condition on jump height (p = 0.014, η^2^_p_ = 0.020) and rate of descent (p = 0.001, η^2^_p_ = 0.121). Multiple comparison tests found a significant decrease in mean effective jump height for 20% (p = 0.028, d = 0.88) and 30% (p = 0.013, d = 0.85) AEL conditions compared to the baseline (BW_pre_); however, there was no significant difference in effective jump height for 10% (p = 0.146, d = 0.60) AEL condition (Figure 2A). Participants showed mixed individual responses in effective jump height before (BW_pre_) and after (BW_post_) AEL interventions (Figure 2B), such that there were no significant differences in effective jump height between the non-AEL conditions (p = 0.648, d = 0.136). There was a significant decrease in average descent speed in all three AEL conditions (10%: p = 0.013, d = 1.012; 20%: p = 0.005, d = 1.181; 30%: p = 0.001, d = 1.337) compared to the baseline (BW_pre_) from the multiple comparisons (Figure 2C).

**Figure 2.**
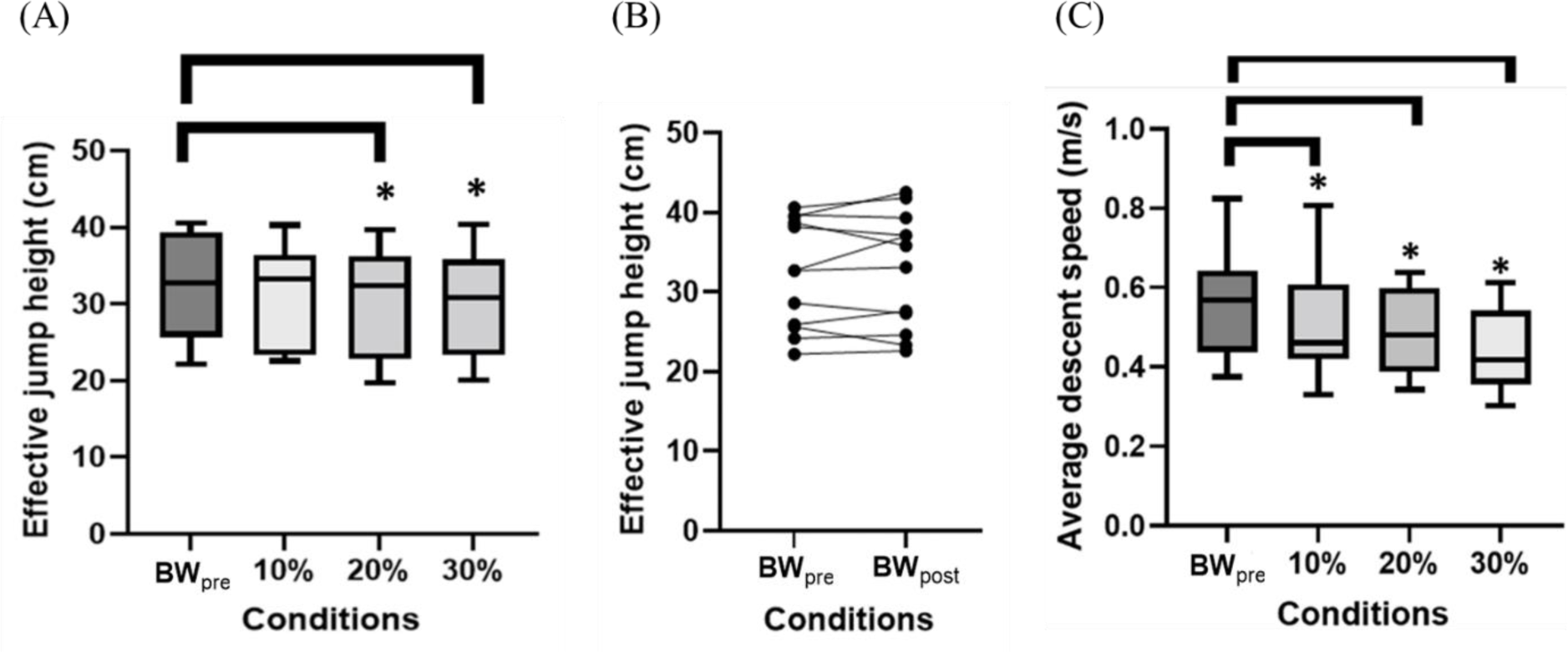
Results for effective jump height and average descent speed. (A) Box and Whisker plot for effective jump height across non-AEL (BW_pre_), 10%, 20%, and 30% AEL conditions. The x-axis shows different loading conditions, and the y-axis shows the effective jump height (cm). The asterisks represent a statistically significant difference from the mean of the baseline (BW_pre_) condition. (B) Effective jump height for non-AEL conditions before (BW_pre_) and after (BW_post_) AEL interventions across individual participants. The x-axis shows different conditions, and the y-axis shows the effective jump height (cm). The black dots represent each selected trial for each participant per condition. (C) Box and Whisker plot for average descent speed across non-AEL (BW_pre_), 10%, 20%, and 30% AEL conditions. The x- axis shows different loading conditions, and the y-axis shows the average descent speed (m/s). The asterisks represent a statistically significant difference from the mean of the baseline (BW_pre_) condition.

### Squat Depth

There was no significant main effect of AEL condition on squat depth (p = 0.107, η^2^_p_ = 0.022). Therefore, we were confident that squat depth control was effectively implemented in this experiment, and there was no systematic bias created by differences in depths across conditions.

### Vertical Ground Reaction Force

The average time-varying VGRF across each condition is shown in Figure 3A. Although VGRF was visually higher with increasing AEL load early in the downward movement, there was no significant difference (p = 0.323, η^2^_p_ = 0.024) in the magnitude of VGRF at the bottom of the countermovement (turning point) across AEL conditions (Figure 3B).

**Figure 3.**
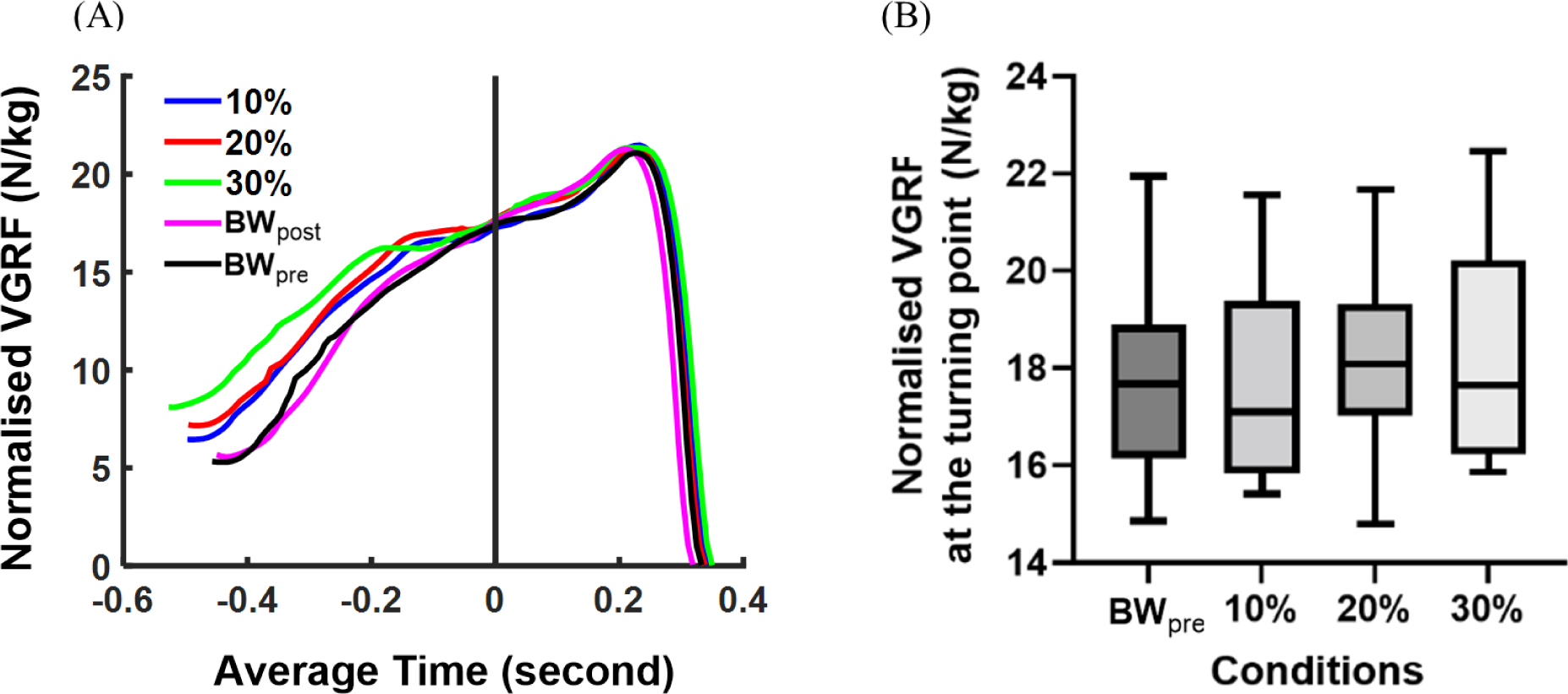
Results for vertical ground reaction force (VGRF). (A) Average time varying VGRF across each condition. The x-axis shows the average time (seconds), and the y-axis shows the body mass normalised VGRF (N/kg). Each colour coded line represents the average value across all participants for each condition. Time 0 represents the bottom of the countermovement (turning point). (B) Box and Whisker plot for body mass normalised VGRF at the turning point across non-AEL (BW_pre_), 10%, 20%, and 30% AEL conditions. The x-axis shows different loading conditions, and the y-axis shows the body mass normalised VGRF (N/kg).

### Joint Kinetics and Energetics

There was a significant main effect of AEL condition on normalised hip extension moments (p = 0.005, η^2^_p_ = 0.107) and knee extension moments (p = 0.014, η^2^_p_ = 0.049) at the turning point. There was a significant decrease in the magnitude of the normalised hip extension moments at the turning point in all three AEL conditions (10%: p = 0.002, d = 1.337; 20%: p = 0.012, d = 1.024; 30%: p = 0.024, d = 0.808) compared to the baseline (BW_pre_) from the multiple comparisons. The normalised knee extension moments at the turning point showed significant increases in 20% (p = 0.005, d = 1.181) and 30% (p = 0.016, d = 1.292) AEL conditions from the multiple comparisons. There were no significant differences across conditions for the normalised ankle plantarflexion moment at the turning point (p = 0.202, η^2^_p_ = 0.030) from the mixed-effect model. The average joint moments (across all joints) relative to time for each condition can be found in Supplementary Figure 3.

The average time-varying right hip, knee, and ankle joint powers across each condition are shown in Figure 4A. There was a significant main effect of AEL condition on positive peak normalised hip joint power during push-off (p = 0.041, η^2^_p_ = 0.059). Multiple comparison tests showed a significant reduction in the positive peak normalised hip joint power during push-off in 10% (p = 0.002, d = 1.371) and 20% (p < 0.0001, d = 1.955) AEL conditions compared to the baseline (BW_pre_). However, the mixed-effect model found no significant differences in positive peak normalised knee (p = 0.215, η^2^_p_ = 0.007) and ankle (p = 0.661, η^2^_p_ = 0.0002) joints power during push-off (Figure 4B).

**Figure 4.**
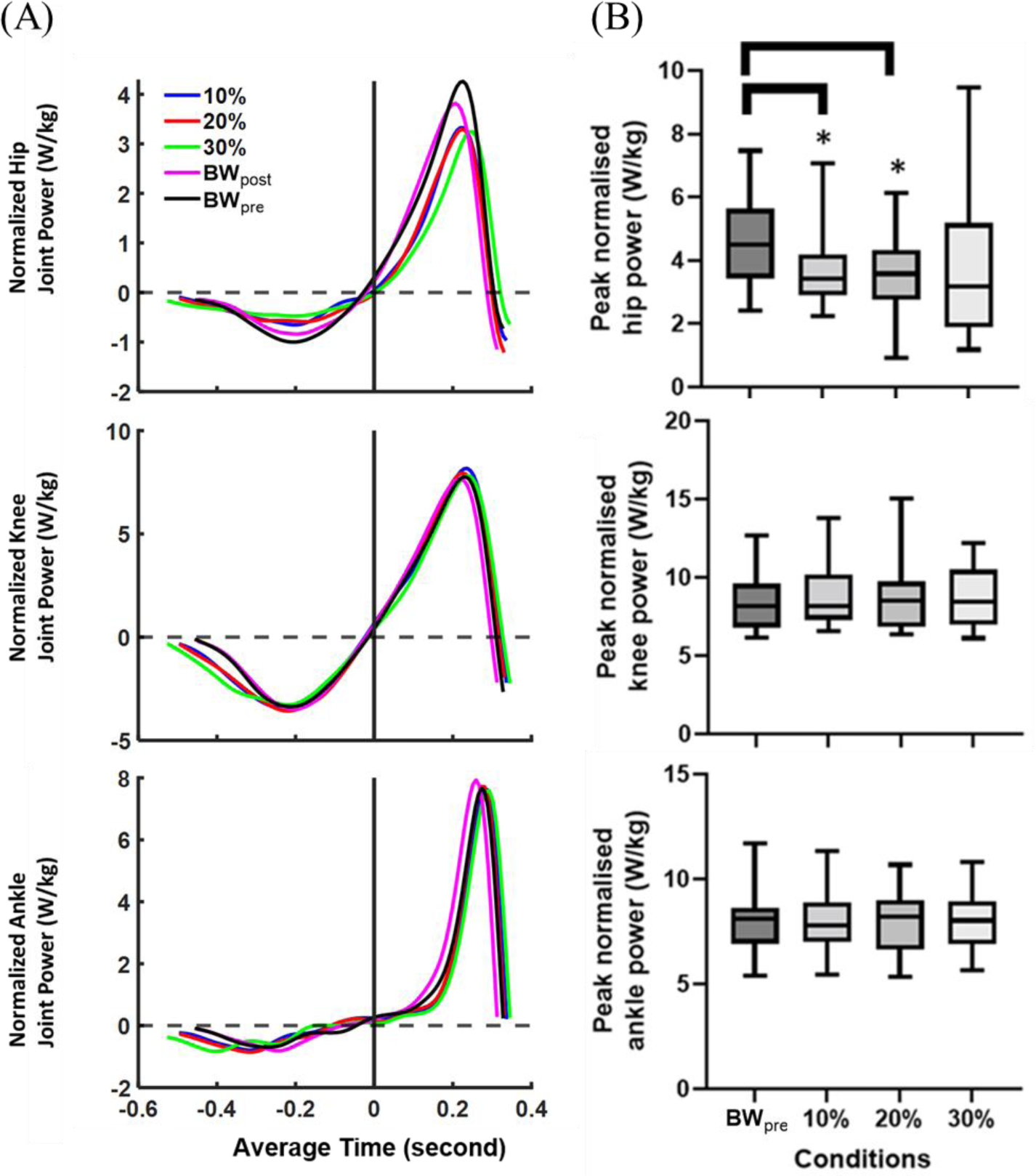
Results for right lower limb joint power. (A) Average time varying right hip, knee, and ankle joint power across each condition. The x-axis shows the average time (second), and the y-axis shows the body mass normalised joint power (W/kg). Each colour coded line represents the average value across all participants for each condition. Time 0 represents the bottom of the countermovement (turning point). (B) Box and Whisker plots for peak normalised right hip extension, knee extension, and ankle plantarflexion joint power during push-off across non-AEL (BW_pre_), 10%, 20%, and 30% AEL conditions. The x-axis shows different loading conditions, and the y-axis shows the peak normalised hip extension, knee extension, and ankle plantarflexion joint power (W/kg). The asterisks represent a statistically significant difference from the mean of the baseline (BW_pre_) condition.

There was a significant main effect of AEL condition on hip joint work (p = 0.001, η^2^_p_ = 0.102) and sum of joints work (p = 0.016, η^2^_p_ = 0.023) during push-off. However, we found no significant main effects in the knee (p = 0.302, η^2^_p_ = 0.016) and ankle (p = 0.418, η^2^_p_ = 0.004) joint work during push-off. Multiple comparison tests showed a significant reduction in the hip joint work during push-off in all three AEL conditions (10%: p = 0.001, d = 1.559; 20%: p = 0.001, d = 1.550; 30%: p = 0.01, d = 0.991) compared to the baseline (BW_pre_). We also found a significant reduction in the sum of joints work in all three AEL conditions (10%: p = 0.005, d = 1.174; 20%: p = 0.017, d = 0.967; 30%: p = 0.001, d = 1.200) compared to the baseline. Data for concentric joint work (across all joints) for each condition can be found in Supplementary Figure 4.

### Vertical COM power and work

The average time-varying concentric vertical COM powers during push-off across each condition are shown in Figure 5A. There were no significant differences (p = 0.207, η^2^_p_ = 0.005) in the magnitude of the concentric peak vertical COM power between AEL conditions (Figure 5B). However, we found a significant main effect of AEL condition on the concentric vertical COM work (p = 0.028, η^2^_p_ = 0.024). Multiple comparison tests showed a significant reduction in the concentric vertical COM work in 20% (p = 0.022, d = 0.926) and 30% (p = 0.005, d= 0.914) AEL conditions compared to the baseline (BW_pre_) (Figure 5C).

**Figure 5.**
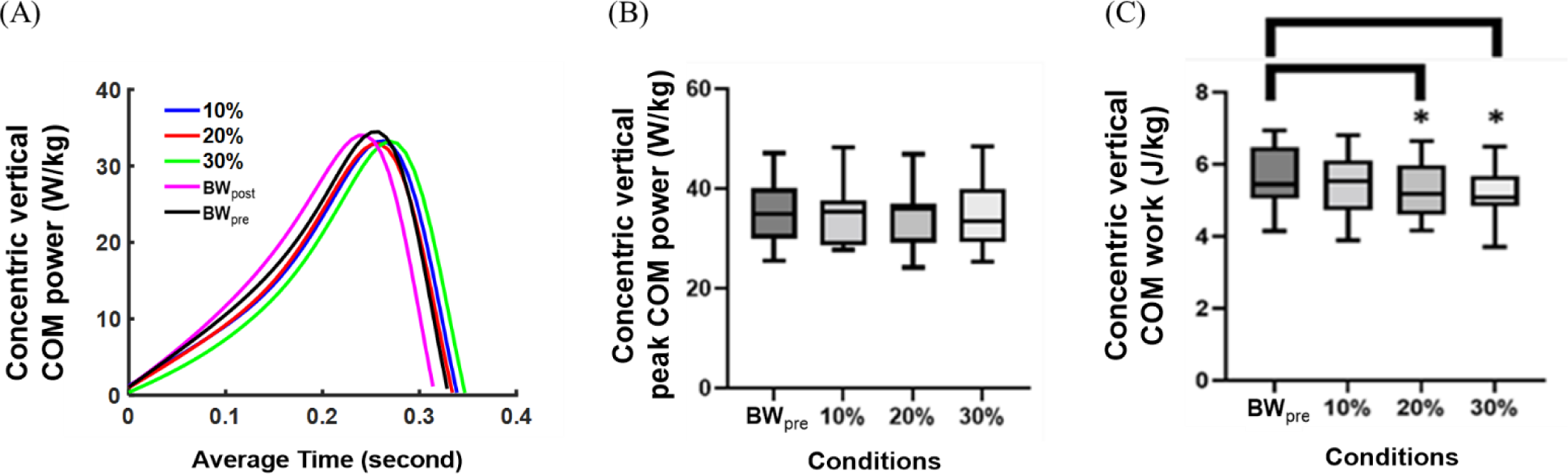
Results for whole-body vertical centre of mass (COM) power and work. (A) Average time varying concentric vertical COM power during push-off across each condition. The x-axis shows the average time (second), and the y-axis shows the body mass normalised COM power (W/kg). Each colour coded line represents the average value across all participants for each condition. (B) Box and Whisker plot for body mass normalised concentric peak vertical COM power across non-AEL (BW_pre_), 10%, 20%, and 30% AEL conditions. The x-axis shows different loading conditions, and the y-axis shows the body mass normalised vertical COM power (W/kg). (C) Box and Whisker plot for concentric vertical COM work across non-AEL (BW_pre_), 10%, 20%, and 30% AEL conditions. The x- axis shows different loading conditions, and the y-axis shows the body mass normalised vertical COM work (J/kg). The asterisks represent a statistically significant difference from the mean of the baseline (BW_pre_) condition.

### Surface EMG

We found no significant differences between conditions in right VL or RF peak EMG amplitudes (VL: p = 0.546, η^2^_p_ = 0.036, RF: p = 0.234, η^2^_p_ = 0.055) or average integrated EMG (VL: p = 0.223, η^2^_p_ = 0.047; RF: p = 0.192, η^2^_p_ = 0.063) during push-off (Figure 6). We also found no significant differences in average peak EMG amplitudes (GLUT: p = 0.502, η^2^_p_ = 0.051, BF: p = 0.785, η^2^_p_ = 0.020, SOL: p = 0.757, η^2^_p_ = 0.020, MG: p = 0.020, η^2^_p_ = 0.172, Supplementary Figure 1) or average integrated EMG (GLUT: p = 0.086, η^2^_p_ = 0.110, BF: p = 0.151, η^2^_p_ = 0.045, SOL: p = 0.312, η^2^_p_ = 0.028, MG: p = 0.135, η^2^_p_ = 0.042, Supplementary Figure 2) between conditions in the other four lower limb muscles, except for the MG muscle which showed a significant reduction in peak EMG (main effect: p = 0.020, η^2^_p_ = 0.172) only in the 20% AEL condition (multiple comparison: p = 0.020, d = 0.957, Supplementary Figure 1).

**Figure 6.**
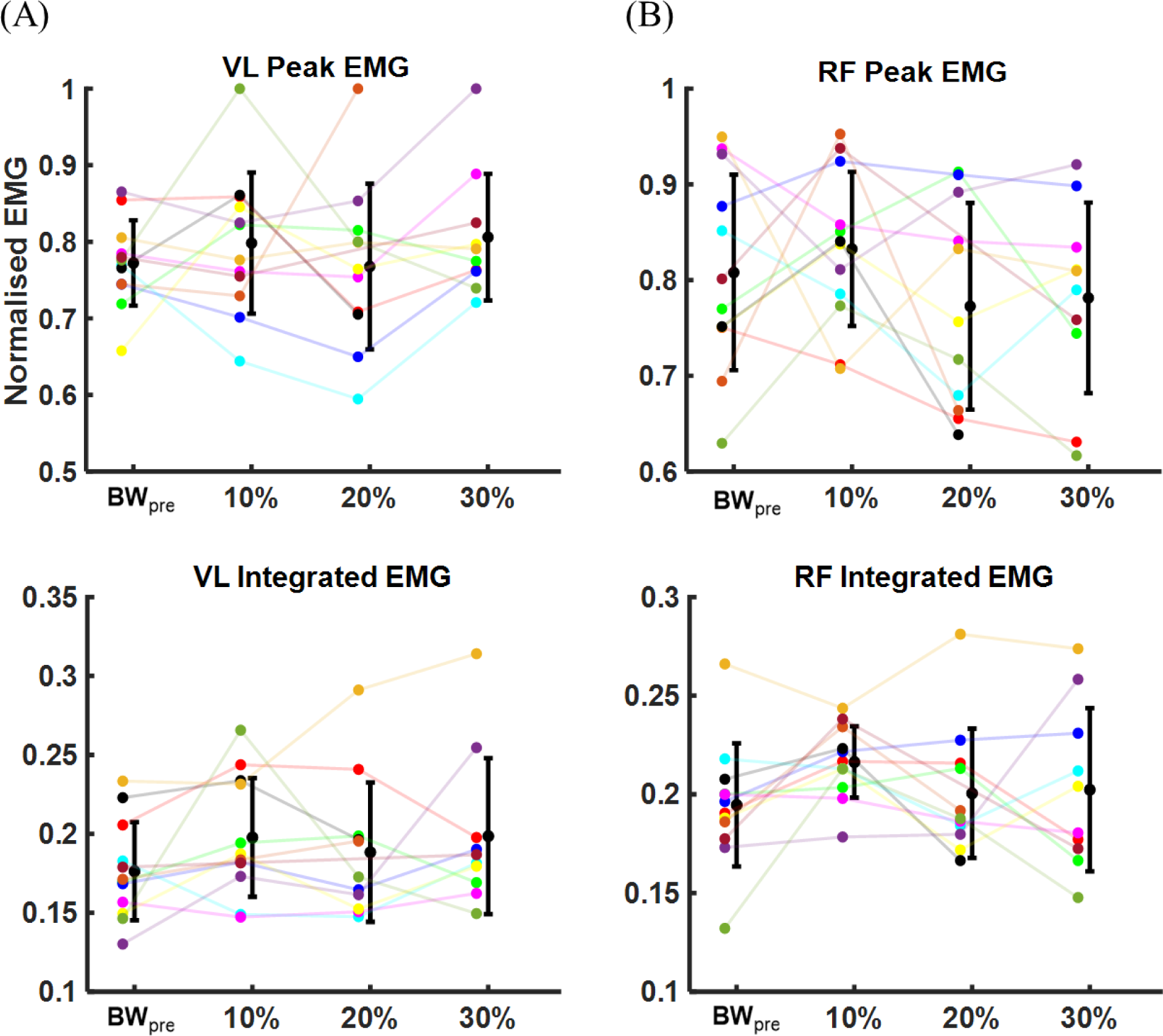
Mean and error bars for the average peak EMG and the average integrated EMG during the entire jump in (A) VL and (B) RF muscles. The x-axis shows different loading conditions, and the y-axis shows the normalised EMG. The smaller dots with different colour codes represent each participant’s average peak or integrated EMG for each condition. The larger black dot represents the group mean, and the error bar represents one standard deviation from the group mean for each condition.

## Discussion

The results show that AEL did not result in performance enhancement in the constrained CMJ used in this study. We found no increase in vertical GRF at the start of the concentric phase nor increases in power or work in the concentric phase. Instead, we found slight decrements in performance in high load conditions. We therefore reject our main hypothesis that added weight alone increases the vertical ground reaction force (VGRF) at the start of the concentric movement and therefore increases the power and subsequent work during the concentric phase of a CMJ. We also reject our second hypothesis because we found no increase in the activation of lower limb muscles during the upward motion with AEL.

Under the strictly constrained conditions imposed here, which controlled contributions from alternative factors that might affect human jump performance (i.e. squat depth, trunk and arm motion), the AEL intervention was found to decrease CMJ performance in the higher added mass conditons. For example, participants showed a decreased COM work during push-off (Figure 5C) and decreased effective jump height (Figure 2A) for 20% and 30% AEL conditions compared to the baseline (BW_pre_). This impaired performance during the AEL jump was associated with a small, but significant reduction in hip joint power (Figure 4A and 4B) and work during push-off, while other joints showed no change in work production.

Given that the post AEL trial jump performance (BW_post_) was unchanged at the group level compared to that before the AEL trials (BW_pre_), the impairments in performance with AEL are unlikely to be a result of order effects or fatigue. Aboodarda et al. ^1^ and Sheppard et al. ^2^ reported that AEL improved CMJ jump height and other concentric kinetic and kinematic parameters (i.e., peak COM power, peak VGRF, and peak COM velocity). However, Aboodarda et al. ^1^ reported that participants increased squat depth to accumulate a longer contact time and larger impulse during push-off to improve jump height. While Sheppard et al. ^2^ did not observe any significant change in squat depth with AEL, neither the joint nor segment kinematics were reported. Potentially, participants in Sheppard et al. ^2^ might have selected different body configurations at the start of the push-off phase which helped to achieve superior performance. Changing body configuration, particularly the trunk angle at the bottom of the countermovement jump, could impact maximal jump performance. ^19^ Our study strictly controlled squat depth and body configuration at the bottom of the countermovement, which eliminated these possible effects during constrained AEL jump.

Aboodarda et al. ^1^ reported an increased rate of eccentric loading and descent velocity during AEL, which could have contributed to performance enhancement via a more rapid stretch-shortening cycle. Previous research shows that applying a faster rate of muscle stretch improves muscle work and power more than a slower stretch. ^20,21^ In contrast, a study that involved performing barbell squats with a jump did not gain performance benefit by adding AEL during the eccentric phase of the squat. ^7^ Moore et al. ^7^ suggested that the relatively slow descent speed during the AEL barbell squat could have limited the performance enhancement effect of AEL. Our present study did not control the rate at which the descent phase was performed, and the slower descent speed observed during heavier AEL conditions might explain why our participants showed decreased performance compared to the non-AEL condition (Figure 2C). An alternative explanation could be that Aboodarda et al. ^1^ used elastic resistance instead of additional mass, which potentially changed the neuromuscular responses due to different loading mechanisms. On the other hand, the task of Moore et al. ^7^ required balancing a barbell and hence was more restrictive in trunk/hip flexion during the eccentric phase. As such, Moore’s et al. ^7^ task required a movement pattern similar to our study, which may explain the similar findings (i.e. no AEL enhancement).

Our results question whether AEL performance effects can be attributed to enhanced ability to store and return energy with additional loading in the eccentric phase. In this study, squat depth and trunk rotation were purposefully constrained to examine the effect of elastic energy return in a strictly controlled environment. Theoretically, if increased elastic energy storage was the major contributing mechanism for AEL’s performance-enhancing effect, we should have observed increased force and joint moments (i.e. VGRF, muscle-tendon unit force) and hence increased elastic energy stored in the tendon under these conditions. Whilst VGRF was higher through most of the downward movement with greater added mass (Figure 3A), the VGRF at the bottom of the countermovement was similar across all conditions (Figure 3B).

We also found that the key joints for storage and return of energy (ankle and knee) showed no or trivial change in joint moment (and hence muscle-tendon force) at the bottom of the countermovement. There was no significant change in ankle joint moment at the turning point across conditions. The increase in knee joint moment at the turning point was significant for 20% and 30% AEL conditions, but it was only 10.5% and 8.2% greater than the BW_pre_ condition. Compared to the reduction in hip joint moment (i.e. 29% reduction in 30% AEL condition compared to BW_pre_), the changes in knee joint moment were minor. Our results showed that higher AEL loads resulted in decreased hip joint moment, power, and work.

Since our study limited the hip and trunk motion, it might be that hip and trunk flexion/extension was important in the mechanism of AEL to increase jump height, and that constraining the hip and trunk motion removed the key performance-enhancing component from AEL. Blache and Monteil ^22^ found that erector spinae muscles contributed to maximal jump performance by adding energy through lumbar spine extension. Potentially, participants in Sheppard et al. ^2^ experienced a greater erector spinae loading with hand-held dumbbells, which might have resulted in higher erector spinae muscle activation and added more energy to the jump during push-off. However, this speculation requires further investigation.

The results of this study support our earlier simulation study which found no performance enhancement in AEL for a simplified single-joint model. ^23^ Our present study showed that added mass did not increase the VGRF at the bottom of the countermovement despite a greater mean force during the desent (Figure 3A). Mechanically, this was achieved by generating force earlier in the descent and increasing the duration of the descent (Figure 3A), which inevitably slowed down the average descent speed in AEL conditions (Figure 2C). This movement characteristic was also predicted in our simulation study. ^23^ Our findings suggest that increased tendon-loading and elastic energy return is unlikely to contribute to the change in performance in a constrained human jumping motion. It is likely that other potential mechanisms, such as increasing squat depth, rate of descent or altered kinematics might be more relevant and contribute to enhanced performance.

There are some important limitations of our research that should be discussed. First, our study used a constrained jump in order to eliminate possible confounding effects during the experiment (i.e. change in squat depth or body configuration). While this was an important experimental control that added strength to our study, we could not directly compare our results to an unconstrained condition in which AEL had a performance-enhancing effect. The statistical power of our study was set to detect changes measured during unconstrained jumps. Whilst it is possible that we were statistically underpowered given the changes in the constraints, the significant reduction in jump height and power measured with AEL would suggest that increasing participant numbers is unlikely to change the result. Another limitation was that we examined recreationally active adults, whereas previous studies typically used trained athletes with significant jumping experience and performance. ^1–4^ It is possible that untrained jumpers are not able to adapt to changes in mass as quickly as trained jumpers. Finally, although we used an objective method to release the mass at the beginning of the upwards movement, it is likely that there were small differences in the release timing across trials that might have contributed to the variability in the jump performances. In our experiment, the average release timing was 0.084 seconds after the turning point, with a small standard deviation of 0.05 seconds. In previous studies, participants were able to release weights at their own discretion, ^2,4^ which probably also causes variation in release times relative to the movement. Future studies might specifically examine the effect of release timing on performance outcomes.

## Conclusion

In this study, AEL did not enhance performance during a trunk-constrained, knee-dominant maximal jump. We found that adding loads to the body did not change the knee extensor muscles’ peak or integrated activation level, nor did it change the maximal VGRF at the start of the push-off. Therefore, AEL did not effectively increase force generation during the push-off phase after the load was released. Furthermore, AEL reduced the effective jump height in 20% and 30% AEL conditions, primarily due to a reduction in hip joint work. As such, we reject the premise that AEL increases jump performance due purely to increased muscle tension and hence storage of elastic energy. It is likely that other mechanisms related to rate of descent, squat depth, or body configuration changes in response to added weight contribute more to any potential AEL effects on jump performance.

## Acknowledgements

We are grateful to Prof. Andrew Cresswell (The University of Queensland) for assistance in developing custom Spike2 scripts in this study.

## Data availability

All relevant data can be found within the article and its supplementary information.

## Authors’ Contribution

Mr. Eric Yung-Sheng Su conceived and designed the experiments, performed the experiments, analysed the data, prepared figures, authored and reviewed drafts of the article, and approved the final draft.

Prof. Timothy J. Carroll conceived and designed the experiments, analysed the data, authored and reviewed drafts of the article, and approved the final draft.

A/Prof. Dominic J. Farris conceived and designed the experiments, analysed the data, authored and reviewed drafts of the article, and approved the final draft.

Prof. Glen A. Lichtwark conceived and designed the experiments, analysed the data, authored or reviewed drafts of the article, and approved the final draft.

